# Interplay between pollution and avian influenza virus in shorebirds and waterfowl

**DOI:** 10.1101/2023.02.23.529812

**Authors:** Tobias A. Ross, Junjie Zhang, Michelle Wille, Alexandros G. Asimakopoulos, Veerle L. B. Jaspers, Marcel Klaassen

## Abstract

Anthropogenic pollution may disrupt wildlife immune function and increase susceptibility to, and ability to withstand, infection. Of particular concern is avian influenza virus (AIV), which in its low-pathogenic form is endemic in many wild bird populations, notably waterfowl and shorebirds, and in its high-pathogenic form poses a threat to wildlife, livestock and people. Many pollutants have immunomodulative properties, yet little is known about how these pollutants affect AIV infection risk specifically. We examined concentrations of known immunomodulatory compounds, per- and polyfluoroalkyl substances (PFASs), and assessed their influence on AIV infection in three reservoir species, red-necked stint (*Calidris ruficollis*, n=121), pacific black duck (*Anas superciliosa*, n=57) and grey teal (*Anas gracilis*, n= 62). Using data on viral prevalence (cloacal/oropharyngeal swabs) and seroprevalence (AIV anti-nucleoprotein antibodies), we found no significant effect of PFASs pollution (total PFASs <0.01 – 470 ng/g in red-necked stint, <0.01 – 600 ng/g in pacific black duck and 0.3 – 200 ng/g in grey teal) on infection status in our three species. This may be due to relatively low pollutant concentrations, but we cannot rule out possible population culling through a synergy of pollution and infection stressors. We therefore recommend further studies on infection incidence in more polluted populations or species.

## Introduction

Pollution threatens wildlife species all around the globe through a diversity of effects. At high enough concentrations, chemical pollution can be acutely toxic, causing direct mortality (Katagi & Fujisawa, 2021). At sub-lethal levels, pollutants may cause physiological harm both individually (Dietz et al., 2019), and when interacting with one another. These interactions can result in other additive or even synergistic harmful effects (Dennis et al., 2020). Physiological effects of pollutants are diverse and may include disruption of both endocrine systems (Dietz et al., 2019; Sebastiano et al., 2021) and reproductive function (Dietz et al., 2019), an increase in oxidative stress (Valavanidis, Vlahogianni, Dassenakis, & Scoullos, 2006), and immunomodulation (Castaño-Ortiz, Jaspers, & Waugh, 2019; Vallverdú-Coll, Mateo, Mougeot, & Ortiz-Santaliestra, 2019). Immunomodulation can change the way organisms interact with pathogens in their environment, thus impacting individuals’ susceptibility to and recovery from disease with knock-on effects on the epidemiology of pathogens.

Interactions between pollution and pathogens can be complex. Direct effects have been studied in a range of pollutants; lead, for example, increases susceptibility to infections by altering T-lymphocyte response in domestic chickens (*Gallus gallus*) (Vallverdú-Coll et al., 2019). The same effects have also been reported from air contaminants from oil sands (including ozone, sulphur dioxide, and polycyclic aromatic hydrocarbons) in wild tree swallows (*Tachycineta bicolor*) (Cruz-Martinez et al., 2015). Mercury pollution has been shown to cause a reduction in natural antibodies in barnacle goslings (*Branta leucopsis*) (Han, van den Berg, Loonen, Mateo, & van den Brink, 2022). Expression of different immune genes in embryonic chicken fibroblasts can be reduced by both polychlorinated biphenyls (PCBs: Badry, Jaspers, & Waugh, 2020) and per-/poly-fluoroalkyl substances (PFASs: Castaño-Ortiz et al., 2019). Indirect interactions with immune function can also emerge with pollution potentially reducing an animal’s body condition resulting in limited availability of resources to mount an immune response to combat infection (Teitelbaum et al., 2022). While immunomodulatory effects of many pollutants have been well studied *in vitro* and in captive birds, studies of both pollution and disease simultaneously in wild birds have rarely been conducted (Teitelbaum et al., 2022).

PFASs are a growing immunomodulatory contaminant group that consist of partially or fully fluorinated carbon chains. This structure lends them both hydro- and lipophobic properties and high stability (Buck et al., 2011). These properties have led to the use of PFASs in many products, including firefighting foams, textiles and kitchen utensils. PFASs are rapidly increasing in diversity with some estimates suggesting over 14,000 are currently in circulation (US Environmental Protection Agency, 2022) with little regulation in place (UNEP, 2022). The immunomodulative properties of PFASs can affect both adaptive and innate immunity of wildlife. In birds, Peden-Adams et al. (2008) showed a reduction of the adaptive response in response to the phytohemagglutinin (PHA) test in chickens (*Gallus gallus domesticus*) exposed to PFOS while in the egg, whereas Smits and Nain (2013) showed a significant reduction of the T-cell dependent antibody response in Japanese quails (*Coturnix japonica*). Effects on innate immunity in chickens include a downregulation in the expression of immune genes nuclear factor kappa-light-chain-enhancer of activated B cells (*NF-κB)*, interleukins 8 (*IL-8*) and 4 (*IL-4*), all of which are involved in the inflammation response (Castaño-Ortiz et al., 2019). Migratory birds of the East-Asian Australasian flyway (EAAF) spending the non-breeding period in Australia may experience exposure to low pathogenic avian influenza virus (AIV), but are potentially also exposed to highly pathogenic avian influenza while in Asia (Wille et al., 2019). Highly pathogenic avian influenza causes severe morbidity and mortality in wild birds, and is currently causing a panzootic with profound impacts on birds, globally (Wille & Barr, 2022). AIV is endemic in Australian wild birds (Wille et al., 2023), and is often associated with other co-infections (Wille et al., 2018), and may thus function as an indicator of overall susceptibility to infection.

In this study, we investigated the effect of PFAS pollution on the infection risk of AIV in three of the pathogen’s prime Australian reservoir species. We focussed on three avian species: one species of shorebird, red necked stint (*Calidris ruficollis*, order Charadriiformes), and two waterfowl species (order Anseriformes): pacific black duck (*Anas superciliosa*) and grey teal (*Anas gracilis*). Long-term AIV prevalence and AIV seroprevalence levels in these three species were established at 3% and 10%, 7% and 50%, and 6% and 55%, respectively, which are among the highest AIV prevalence values measured in birds in Australia (Wille et al., 2023). Furthermore, evidence of highly pathogenic avian influenza infection has previously been found in red-necked stint (Wille et al., 2019). This species also migrates via an extremely polluted region (including PFASs), China’s Yellow Sea (Muir & Miaz, 2021; Xiao et al., 2017), highlighting the potential threat posed by the interplay between pollution and disease. As PFASs (and PFOS in particular), have been shown to downregulate the innate immune response (Castaño-Ortiz et al., 2019), we hypothesised that with increasing exposure to these pollutants the incidence of both AIV and AIV antibodies would increase.

## Methods

### Ethics statement

All birds were captured and sampled under approval of Deakin University Animal Ethics Committee (permit numbers A113-2010, B37-2013, B43-2016, B39-2019), Wildlife Ethics Committee of South Australia (2011/1, 2012/35, 2013/11), and Philip Island Nature Park Animal Ethics Committee (SPFL20082).

### Sample collection

Red-necked stint were captured on the coast of Victoria, at the Western Treatment Plant (WTP) and in Western Port Bay, both near Melbourne, Australia. Pacific black ducks were sampled at the WTP, and on farmland near Geelong, also in Victoria. Grey teal were also sampled at the WTP and Innamincka Regional Reserve in South Australia. Catches of birds occurred from 2011-2020. Red-necked stint were captured using cannon-nets in collaboration with the Victorian Wader Study Group (VWSG), as part of ongoing shorebird banding work starting 1978 (Minton, 2006). Ducks were caught using baited walk-in traps and mist nets. All birds were blood sampled and oropharyngeal and cloacal swabs were taken. Of each bird, cloacal and oropharyngeal swab samples were collected using sterile swabs, and immediately stored in virus transport medium (VTM, brain heart infusion [BHI] broth-based medium [Oxoid] with 0.3 mg/ml penicillin, 5 mg/ml streptomycin, 0.1 mg/ml gentamicin, and 2.5 g/ml, amphotericin B), either separately (pre-March 2014) or together (March 2014 onwards). Swab samples in VTM were kept refrigerated for up to a week, before being stored at below -80°C. Blood samples (typically 200 μl) were collected from the brachial vein of each bird using Sarstedt Microvette capillary tubes. Samples were refrigerated and left to clot, next centrifuged within 6-24 hrs after collection to separate red blood cells (RBC) from serum, after which they were stored at below -20°C. Serum was used to test for the presence of AIV antibodies, while RBC was used for PFAS analyses.

Across the entire influenza surveillance study, we collected 1512 sample sets from red necked stints, 446 from pacific black ducks, and 660 from grey teals. We selected the samples of all 41 AIV-positive red-necked stints, 29 AIV-positive pacific black ducks and 29 AIV-positive grey teals. These samples were matched with the samples from 68, 28 and 33 AIV negative birds, respectively. The AIV negative sample sets were stratified-randomly selected, ensuring at least 10 AIV negative individuals were included from each of the sites where AIV positive birds were sampled. Serum samples from the same birds were also used for AIV antibody analyses with the exception of two pacific black ducks and three grey teals for which we failed to collect serum. For red-necked stint we included an additional 12 serum samples for serological analysis. Total numbers of individuals involved across all samples were 121 red-necked stints, 57 pacific black ducks and 62 grey teals.

### AIV screening

Samples were screened for AIV following established methods, described in Wille et al. (2023). Briefly, RNA was extracted from swab samples, which were assayed for AIV using quantitative reverse transcriptase real time PCR targeting a short fragment of the matrix gene (Spackman et al., 2002). A cycle threshold (Ct) cut-off of 40 was used. In cases where oropharyngeal and cloacal swabs were collected separately (pre-March 2014), results were combined and a bird was deemed AIV-positive if either swab returned a positive result. Blood serum samples were assayed for anti-nucleoprotein antibodies of AIV using the Multi Screen Avian Influenza Virus Antibody Test Kit (IDEXX, Hoppendorf, The Netherlands) following manufacturer’s instructions.

### Chemical analyses of blood samples

All RBC samples were analysed for 12 different PFASs (full details provided in Table S1), which were broadly grouped as carboxylates (PFPA, PFHxA, PFOA, PFNA, PFDA, PFUnA, PFDoA, PFTrA, PFTeA) and sulfonates (PFBS, PFOS). We also calculated a total PFAS group, consisting of all carboxylates and sulfonates listed above, as well as PFOSA. Analyses were conducted using methodology adapted from Trimmel et al. (2021). In brief, ∼ 50 mg of red blood cell samples was weighted in a 1.5 mL polypropylene tube and 10 μL of PFOA-^13^C_8_ and PFOS-^13^C_8_ (1000 μg/L) was added. After adding 0.3 mL methanol containing 1% ammonium formate, the sample was vortexed for 30 s and ultrasonicated for 30 min. Then the sample was centrifuged for 5 min at 3500 rpm. Finally, the supernatant was transferred into a 1.5 mL injection vial with an insert vial after purification by Hybrid-SPE. The sample extract was stored at − 20 °C until analysis by UPLC-MS/MS.

### Statistical analysis

All statistical analyses were conducted using R Version 4.2.0, in RStudio 2022.02.0 Build 443. All PFASs were summed into carboxylate (ΣPFCAs), sulfonate (ΣPFSA), and total PFASs (ΣPFAS) categories as outlined above. All non-detected values of individual compounds were replaced with 0.015 ng/g, equivalent to half the minimum quantification limit of detection for all compounds. When summed categories were calculated, any totals where no compounds were detected were also set at 0.015 ng/g. For each of the three focal species, we first ran two-sample t-tests on the grouped PFAS data (ΣPFCAs, ΣPFSAs and ΣPFASs) from each region, to determine if there were pollution differences between regions.

We used generalised linear mixed effects models in R package lme4 (Bates, Mächler, Bolker, & Walker, 2015) (function glmer, family binomial) to investigate AIV status (positive or negative) in each species, as a function of each of our 12 pollutants individually, as well as ΣPFCAs, ΣPFSAs, and ΣPFASs (i.e. 3 × 15 = 45 models). These models included region as a random effect. We also ran this same structure of models across all samples, including species as a second random effect alongside region (i.e. 15 models). Finally, we ran the same model across all samples combined (again with species and region as random effects), in which ΣPFCAs and ΣPFSA concentrations were both entered simultaneously as explanatory variables (i.e. 4 models). Overall, we thus ran a total of 64 models for AIV prevalence. We ran the same set of models using AIV serostatus (positive or negative) instead of AIV status as dependent variable, culminating in a grand total of 128 models testing the effect of PFASs on AIV status and serostatus.

## Results

Of the 12 individual pollutants that we targeted, only PFOS, PFDoA and PFUnA were detected in more than 50% of our samples (94%, 72% and 60% respectively, Tables 1 and 2). When summed into overall categories, ΣPFCAs, ΣPFSAs and ΣPFASs were detected in 90%, 95% and 98% of all samples respectively, regardless of species. Across all the 240 samples tested, ΣPFCAs were not detected in 24 samples, ΣPFSAs were not detected in 13 samples, and ΣPFASs were not detected in only six samples. ΣPFCAs concentrations were below 50 ng/g in all but 7 samples (3%), with samples from four red-necked stints (56, 71, 93 and 98 ng/g) and three pacific black ducks (54, 70 and 74 ng/g) comprising the exception. ΣPFSAs by contrast were higher in concentration, with 11 samples (5%) exceeding 200 ng/g (Figure 1), comprising of 6 red-necked stints (223-423 ng/g) and 5 pacific black ducks (208-503ng/g). We found that all ΣPFCAs, ΣPFSAs and ΣPFASs were significantly higher at the WTP in all species (*p*<0.05, see Table S2 for summary concentrations), with one exception where no significant differences between regions was found for ΣPFCAs in pacific black duck. A complete range of detection rates and overall concentrations by AIV status and AIV serostatus are shown in Table 1 and Table 2, respectively.

**Table 1.**
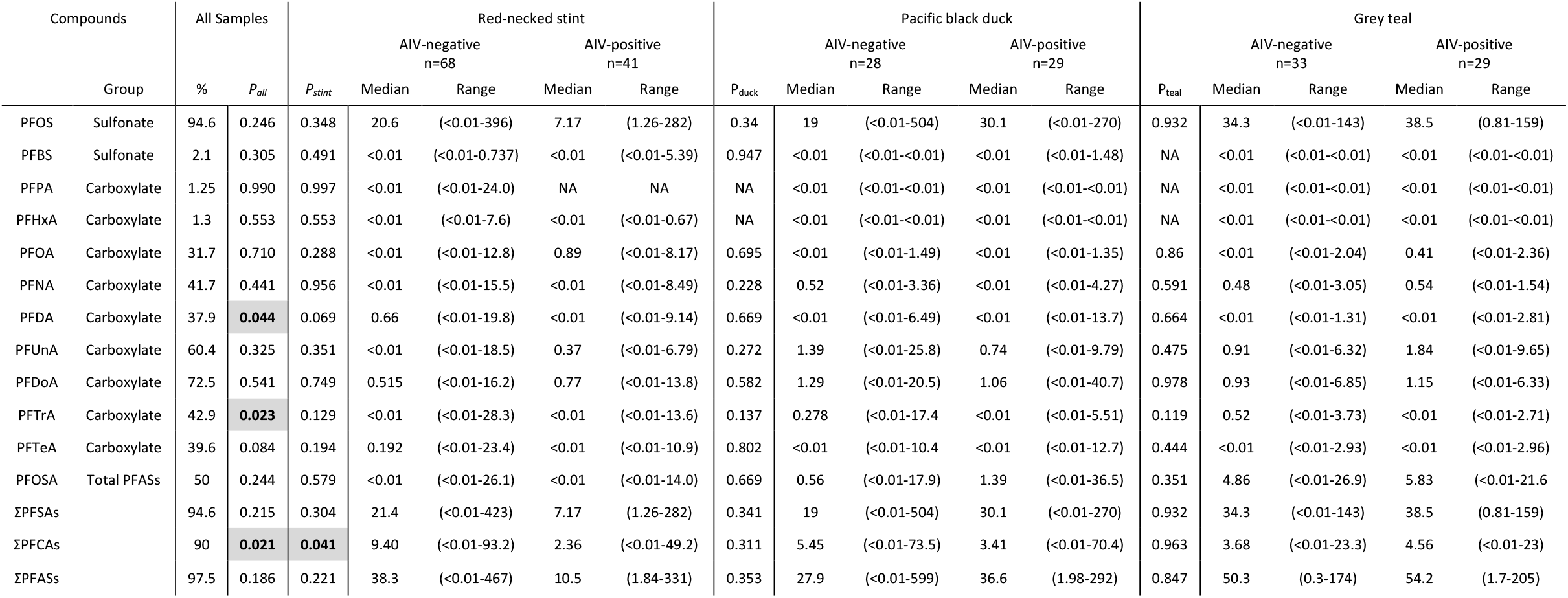
Median and range concentrations of PFASs (in ng/g) in red blood cells of red-necked stint (*Calidris ruficollis*), pacific black duck (*Anas superciliosa*) and grey teal (*Anas gracilis*), based on the AIV infection status of each individual. Groups of compounds have been indicated. Where <0.01 is noted, this means the concentration was below the limit of quantification. Emboldened and shaded *p-*values indicate statistical significance between AIV negative and positive individuals. Where a value is recorded as NA, this compound was not detected in the species.

**Table 2:**
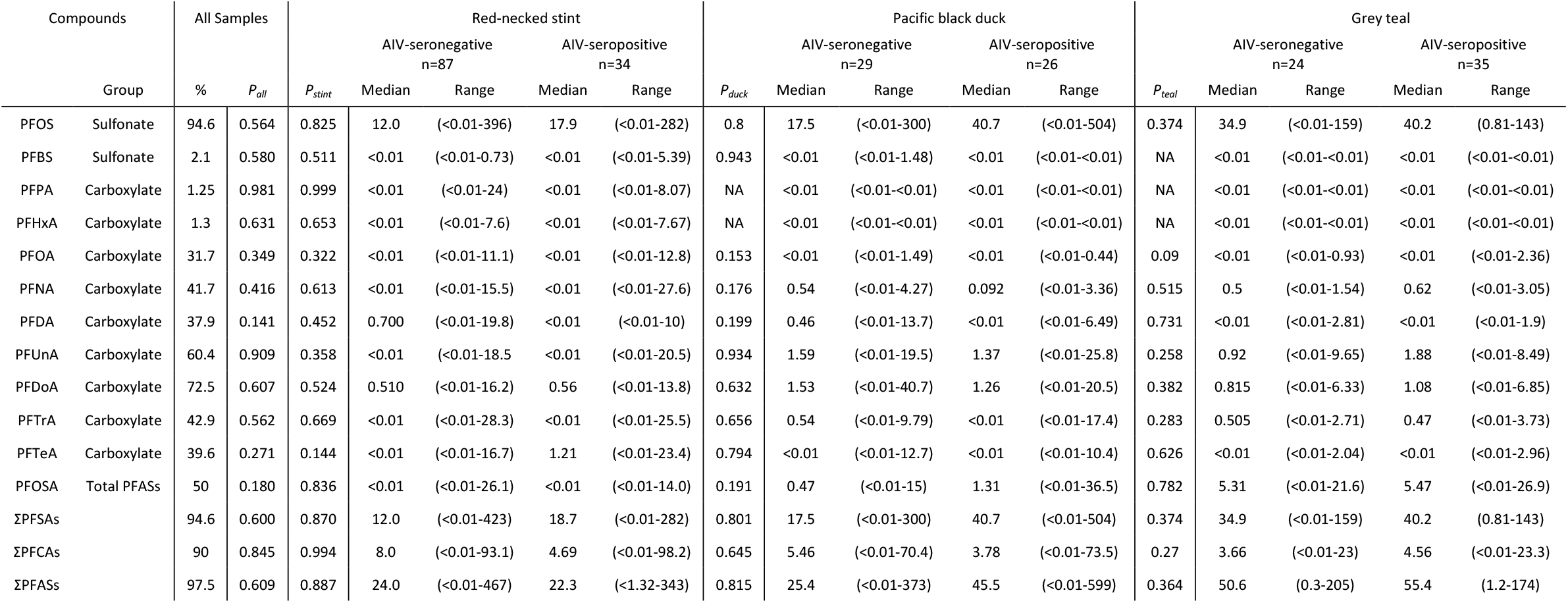
Median and range concentrations of PFASs in red blood cells (in ng/g) of red-necked stint (*Calidris ruficollis*), pacific black duck (*Anas superciliosa*) and grey teal (*Anas gracilis*), based on AIV serostatus of each individual. Groups of compounds have been indicated. Where <0.01 is noted, this means the concentration was below the limit of quantification. Emboldened and shaded *p-*values indicate statistical significance between seropositive and seronegative individuals. Where a value is recorded as NA, this compound was not detected in the species.

**Figure 1:**
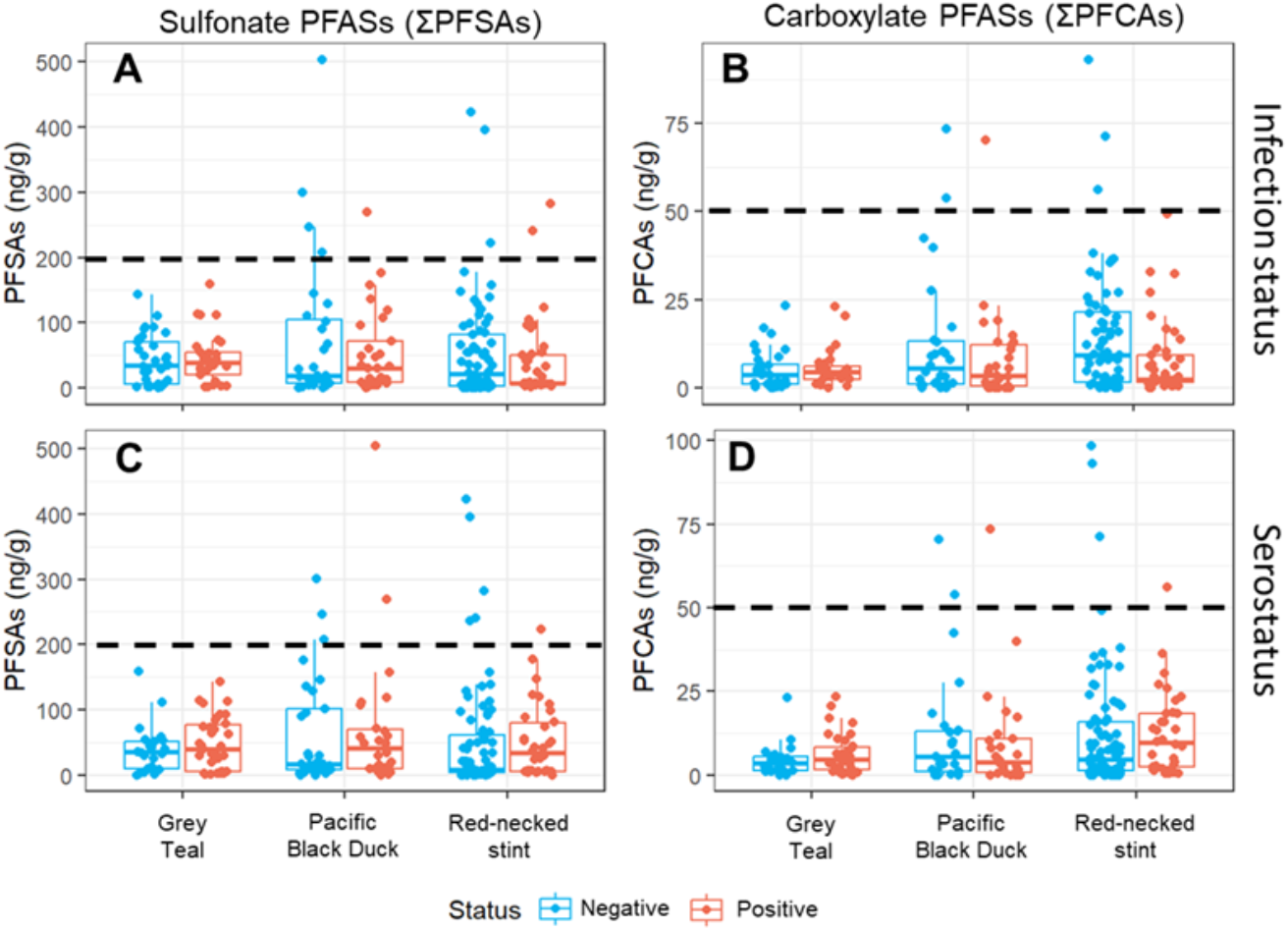
Concentrations (ng/g) of summed sulfonate (A,C) PFASs (ΣPFSAs) and summed carboxylate (B, D) PFASs (ΣPFCAs) in red blood cells from grey teal (*Anas gracilis, n*=62), pacific black duck (*Anas superciliosa*, n=57) and red-necked stint (*Calidris ruficollis*, n=121). A and B show infection status, while C and D show serostatus. Birds that were positive are represented in red, and negative in blue. 4.5% of samples exceeded 200ng/g (dashed line in A and C) of ΣPFSAs, while 3.9% of samples exceeded 50ng/g (dashed line in B and D) of ΣPFCAs (dashed line in B and D).

Across all 128 models, no significant effects were found for the influence of PFAS pollution on AIV prevalence and seroprevalence, except for five models (*p* values for each species, and overall sample population provided in Tables 1 and 2). Significant effects were found for PFDA, PFTrA and ΣPFCAs on AIV infection status for all species combined, and for ΣPFCAs on AIV infection status for red-necked stints when modelled individually. Finally, ΣPFCAs were also found to have a significant effect on AIV infection status of red-necked stints when ΣPFSAs were also included in the same model. No significant effect of pollution on serostatus was found in any model. PFDA was found to be lower in birds that were infected with AIV (AIV positive median: <0.01 ng/g, range: <0.01-13.71 ng/g; AIV negative median: <0.01 ng/g, range: <0.01-19.8 ng/g; slope: −0.146, *p* = 0.044), as was PFTrA (AIV positive median: <0.01 ng/g, range: <0.01-13.6 ng/g; AIV negative median: <0.01 ng/g, range: <0.01-28.3 ng/g; slope = −0.129, *p* = 0.023) The same pattern was observed for all three models in which ΣPFCAs were found to have a significant effect, across all species modelled together (AIV positive median: 3.44 ng/g, range: <0.01-98.2 ng/g; AIV negative median: 5.25 ng/g, range: <0.01-93.2 ng/g; slope = -0.028, *p* = 0.021), and in red-necked stints when ΣPFCAs were modelled individually (AIV positive median: 2.36 ng/g, range: <0.01-49.2 ng/g; AIV negative median: 9.40 ng/g, range: <0.01-93.2 ng/g; slope = −0.037, *p* = 0.041) and with ΣPFSAs (slope = −0.048, *p* = 0.049).

## Discussion

Our results, encompassing concentrations of 12 per-/polyfluorinated compounds, and both AIV infection and serostatus in three different species, did not suggest that infection of AIV in these birds is influenced by their PFAS burdens. Of the 128 models run, only five significant correlations were detected with *p* values just below 0.05, which could well be spurious given the number of models that were run. Our general finding of no (or few) significant correlations is in contrast to a previous *in vitro* study where PFASs were demonstrated to have immunomodulatory effects in birds (Castaño-Ortiz et al., 2019). If such effects were present in the current study, we would have expected actively AIV-infected or previously AIV-infected birds to have higher concentrations of PFASs in their system.

We expect that the absence of a correlation between PFAS pollution and AIV in this study was because our observed PFAS concentrations were too low to elicit immunomodulatory effects. In the *in vitro* study by Castaño-Ortiz et al. (2019), exposure to PFOS was at 22mg/L, likely far higher than any exposure our study birds experienced. Indeed, even though PFAS pollution often varied between sample sites, the concentrations we observed were low relative to ‘predicted no-effects concentrations’ in serum presented by Newsted, Jones, Coady, and Giesy (2005), the lowest of which was 150 ng/g. PFAS concentrations in all but three individuals’ (two red-necked stints and one pacific black duck) were also low compared to ‘lowest observable adverse effect levels’ (LOAELs) of 400ng/g in adult bird liver tissues proposed by Dennis et al. (2021). These LOAELs were calculated based on decreased adult weight gain. Unfortunately, the disparity between sample matrices (liver vs. red blood cells) means we cannot draw direct comparisons between our study and Dennis et al. (2021)’s proposed toxicity thresholds. Nevertheless, concentrations of PFASs in liver tend to be higher than those in blood (Gebbink & Letcher, 2012) and as such, we can consider Dennis et al. (2021)’s liver LOAELs to be conservative estimates of toxicity in blood. The true number of our samples that exceed the adult LOAEL from Dennis et al. (2021) is likely to be higher. Aside from the three individual exceptions mentioned above, the low concentrations we observed may be considered remarkable for birds that inhabit putatively polluted wastewater treatment plants (all focal species) or migrate via highly polluted environments (red-necked stint migrating via e.g. the Yellow Sea) (Muir & Miaz, 2021). Nevertheless, in 105 individuals, the total PFASs concentrations found were higher than the lowest LOAELs posited by Dennis et al. (2021) (50 ng/g in liver for young birds, based on decreased successful hatching), which suggested there may be some risk to juvenile birds in our population. The absence of specific data on immunomodulation thresholds means we cannot definitively state whether our concentrations are below levels to cause effect. However, our results did not yet suggest that this is the case.

It is worth noting that some individuals had particularly elevated PFASs concentrations (e.g. one pacific black duck was observed with 599 ng/g of PFOS, with the next highest concentration observed in a red-necked stint at 467 ng/g, see Figure 1), while being negative for both AIV infection. However, no conclusions can be based on such rare extreme cases due to of a) the very small sample size of individuals with elevated PFASs and b) the level of exposure of these pacific black ducks and red-necked stints to AIV (10% seroprevalence: Wille et al., 2023) increasing stochasticity in the occurrence of infection. It is clear though, that individuals can accrue a high PFAS load without necessarily incurring infections.

An alternative explanation for the low PFASs concentrations, and minimal significant effects (aside from 5 out of the 128 models suggesting a negative relationship between AIV infection and pollution burden), is that the absence of an effect is due to population culling. Population culling is the phenomenon whereby sick, or in this case, polluted individuals do not survive and are thereby ‘culled’ from the population, such that we never or infrequently detect them when sampling. In our sample population, there was clearly the possibility for individuals to accrue elevated PFAS burdens, as demonstrated by the 11 birds with PFSAs concentrations greater than 200ng/g (see Figure 1). In the absence of additional stressors such as disease, birds may be able to withstand elevated concentrations. Congruently, wild birds have limited documented signs of disease associated with low pathogenic avian influenza beyond a reduction in daily movement in mallards (*Anas platyrhynchos*) (Van Dijk et al., 2015) and poorer performance in foraging or migrating Bewick’s swans (*Cygnus columbianus*) (Hoye et al., 2016; Van Gils et al., 2007). The limited extent of these signs are such that infection alone has little to no direct effect on the survival of its host (Maxted et al., 2012). However, both the negative effect of pollution in isolation as well as the potential combined effect of pollution and disease might even cause death, such that individuals with downregulated immune responses due to pollution may be unable to survive the combined effects of PFASs and the virus (Guruge et al., 2009). Such individuals may therefore have been lost to our study. Indeed, in all five models where we found significant effects of pollution on AIV infection status, birds with lower pollution were more likely to be AIV positive. The significant model results could indicate that infected birds with higher concentrations are unable to survive the added pressures of both and therefore ‘culled’ from the population. Hence, it is possible that we did not sample highly polluted birds that also had avian influenza infections, because they did not survive. This is particularly pertinent for migrating shorebirds which also deal with the additional stressors of physiological effects of migration including immunomodulation (Buehler, Tieleman, & Piersma, 2010). Indeed, there is some evidence of migratory culling, whereby migration may remove sick individuals (McKay & Hoye, 2016). This effect could be more pronounced under pollution stress, which could become more toxic due to the thermal stress from exertion during migration (Gordon, Johnstone, & Aydin, 2014). This is a possible explanation as to why the only significant single-species models found were models involving data from red-necked stints, as these are the only migratory species included in the study.

Overall, our birds were all sampled in environments with relatively low PFASs concentrations, with the most polluted being processed wastewater at the WTP. In other regions of the world, PFAS pollution may be higher (Muir & Miaz, 2021), as is the prevalence of AIV and therefore the likelihood of a birds’ exposure to the virus (Wille & Barr, 2022). In these regions, the effects of pollution-mediated immunomodulation may be much stronger. The consequences of widespread immunomodulation could be severe, for populations of wild birds, and for human health and poultry industries. Our findings of no clear evidence of PFAS-mediated immunomodulation in Australia cannot discount effects in more heavily polluted regions of the world where exposure to more dangerous high pathogenicity AIV strains is more prevalent. The current global circumstances of increased highly pathogenic avian influenza transmission amongst wild birds in the northern hemisphere highlight the necessity to continue monitoring potential drivers of host susceptibility such as pollution, so that we may garner a better understanding of the threats the interactions of these stressors pose to wild birds.

## Acknowledgements

This project was funded by the Australian Research Council (ARC) Discovery Project Grant DP190101861 (to Marcel Klaassen) and additional funding by the Norwegian Research Council (NRC) project COAST IMPACT 302205/SHJ (to Veerle Jaspers). The WHO Collaborating Centre for Reference and Research on Influenza is funded by the Australian Department of Health. Michelle Wille is funded by an Australia Research Council Discovery Early Career Research Award (DE200100977). We thank the VWSG for field sampling and logistics, in particular acknowledging Rob Patrick, Margaret Bennett, Maureen Christie and the late Clive Minton.

