## Supplementary material for "Interplay between pollution and avian influenza virus in shorebirds and waterfowl": Table S1 (from another version of this preprint: version 1, 2023-02-24)

**SUPPLEMENTAL MATERIALS: Interplay between pollution and avian influenza virus infection in waterbirds**

**Table S1** Chemical names and formulae of PFASs

| **Compound Abbreviation** | **Full name** | **Structure** | **CAS** | **Formula** |
| --- | --- | --- | --- | --- |
| PFPA | Perfluoropentanoic acid | 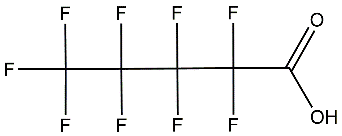 | 2706-90-3 | C_5_HF_9_O_2_ |
| PFHxA | Perfluorohexanoic acid | 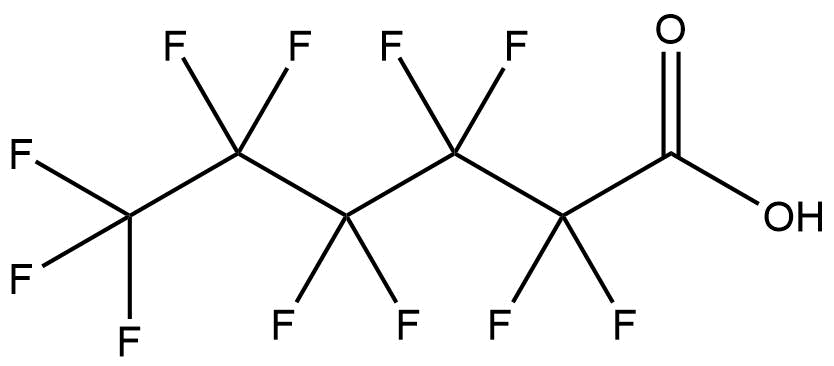 | 307-24-4 | C_6_HF_11_O_2_ |
| PFOA | Perfluorooctanoic acid | 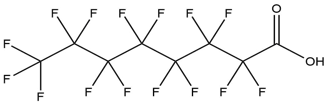 | 335-67-1 | C_8_HF_15_O_2_ |
| PFNA | Perfluorononanoic acid | 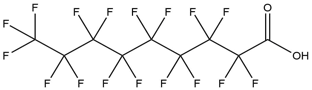 | 375-95-1 | C_9_HF_17_O_2_ |
| PFDA | Perfluorodecanoic acid | 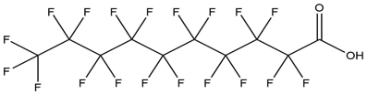 | 335-76-2 | C_10_HF_19_O_2_ |
| PFUnA | Perfluoro-undecanoic acid | 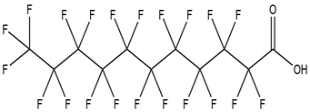 | 2058-94-8 | C_11_HF_21_O_2_ |
| PFDoA | Perfluoro-dodecanoic acid | 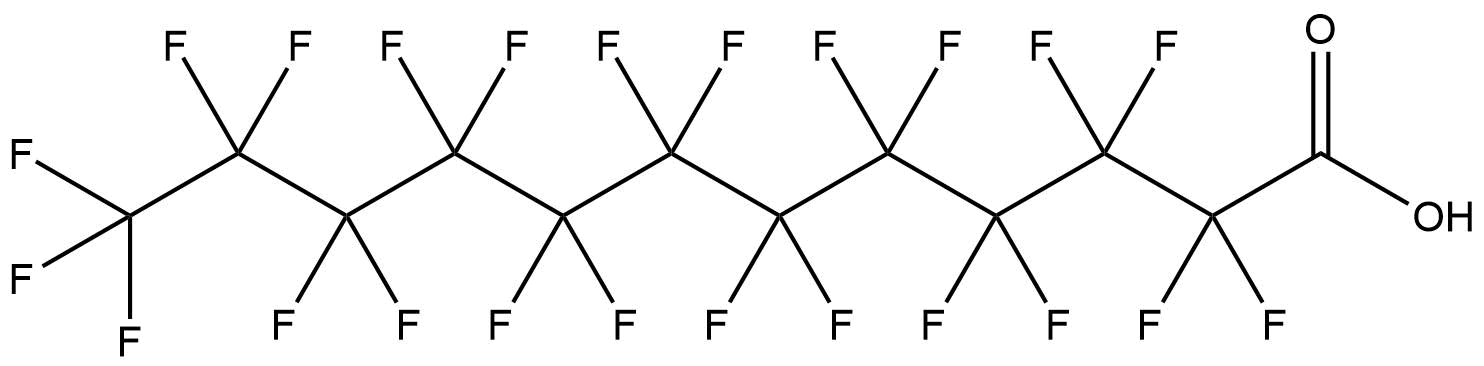 | 307-55-1 | C_12_HF_23_O_2_ |
| PFTrA | Perfluoro-tridecanoic acid | 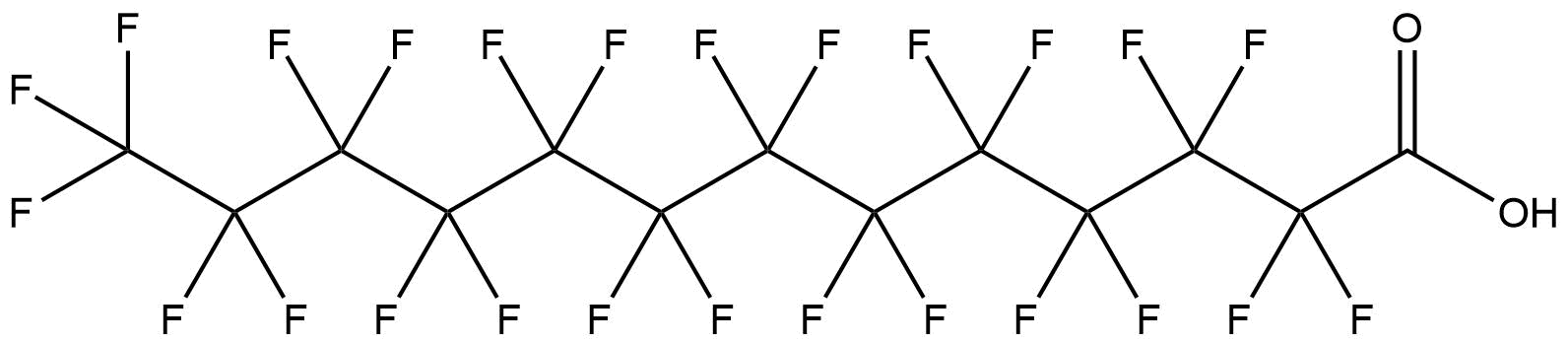 | 72629-94-8 | C_13_HF_25_O_2_ |
| PFTeA | Perfluoro-tetradecanoic acid | 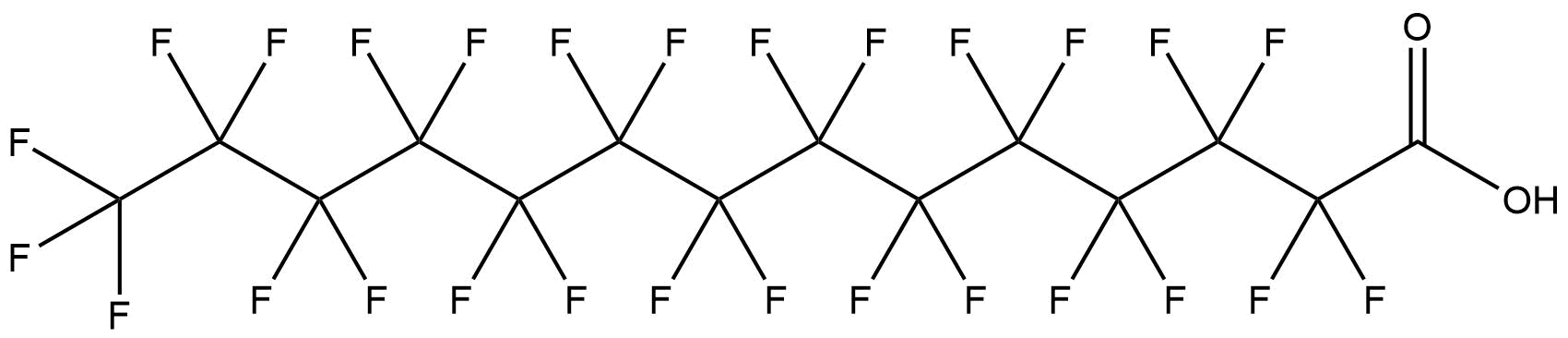 | 376-06-7 | C_14_HF_27_O_2_ |
| PFBS | Nonafluorobutane-sulfonic acid | 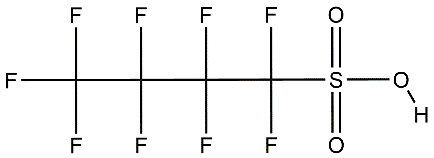 | 375-73-5 | C_4_HF_9_SO_3_ |
| PFOS | Perfluorooctano-sulfonic acid | 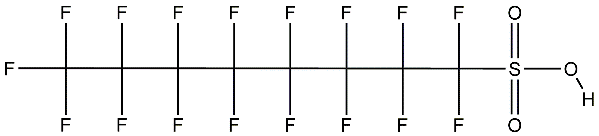 | 1763-23-1 | C_8_HF_17_O_3_S |
| PFOSA | Perfluorooctane sulfonamide | 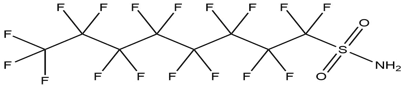 | 754-91-6 | C_8_H_2_F_17_NO_2_S |

**Table S2:** Summed PFAS median and range concentrations for each species at each sample site.

| Site | Compound | Species | | | | | | | | |
| --- | --- | --- | --- | --- | --- | --- | --- | --- | --- | --- |
|  |  | Red-necked Stint | | | Pacific Black Duck | | | Grey Teal | | |
|  |  | n | Median (ng/g) | Range (ng/g) | n | Median (ng/g) | Range (ng/g) | n | Median (ng/g) | Range (ng/g) |
| Western Treatment Plant (WTP), Victoria | ΣPFCAs | 64 | 8.59 | <0.01-98.23 | 33 | 9.12 | <0.01-73.51 | 47 | 4.85 | 0.61-23.3 |
|  | ΣPFSAs |  | 51.13 | <0.01-1275 |  | 71.46 | <0.01-504 |  | 44.82 | 4.87-158.85 |
|  | ΣPFASs |  | 66.7 | <0.01-1293 |  | 101 | <0.01-599 |  | 56.62 | 5.85-205 |
| Western Port Bay (WPB), Victoria | ΣPFCAs | 57 | 1.84 | <0.01-48.9 | -- | -- | -- | -- | -- | -- |
|  | ΣPFSAs |  | 6.67 | <0.01-411 |  | -- | -- |  | -- | -- |
|  | ΣPFASs |  | 9.36 | <0.01-460 |  |  |  |  |  |  |
| Geelong, Victoria | ΣPFCAs | -- | -- | -- | 24 | 1.21 | <0.01-70.37 | -- | -- | -- |
|  | ΣPFSAs |  | -- | -- |  | 10.54 | <0.01-58.89 |  | -- | -- |
|  |  |  |  |  |  | 16.17 | <0.01-70.37 |  |  |  |
| Innamincka, South Australia | ΣPFCAs | -- | -- | -- | -- | -- | -- | 15 | 0.64 | <0.01-7.12 |
|  | ΣPFSAs |  | -- | -- |  | -- | -- |  | 2.31 | <0.01-7.64 |
|  | ΣPFASs |  |  |  |  |  |  |  | 2.83 | 0.3-9.6 |

'--' is used to denote the absence of data as the species was not sampled at this site.
